# Epistasis drives rapid divergence across multiple traits during the adaptive evolution of a carbapenemase

**DOI:** 10.1101/2024.09.10.610996

**Authors:** Laura Dabos, Inssaf Nedjari, Alejandro Couce

## Abstract

Interactions among beneficial mutations (i.e., epistasis) are often strong enough as to direct adaptation through alternative mutational paths. While alternative solutions should display similar fitness under the primary selective conditions, their properties across secondary environments may differ widely. The extent to which these cryptic differences are to be expected is largely unknown, despite their fundamental and practical importance, such as in the search for exploitable collateral sensitivities among antibiotic resistance mutations. Here we use directed evolution to characterize the diversity of mutational paths through which the prevalent carbapenemase KPC-2 can evolve high activity against the clinically-relevant antibiotic ceftazidime, an initially poor substrate. We identified 40 different substitutions, including many common clinical settings, spread along 18 different mutational trajectories. Initial mutations determined four major groups into which the trajectories can be classified, a signature of strong epistasis. Of note, despite minor variation in final ceftazidime resistance, groups diverged markedly across multiple phenotypic dimensions, from molecular traits such as stability and hydrolitic efficiency to macroscopic traits such as growth rate and activity against other β-lactam antibiotics. Our results indicate that cryptic yet consequential phenotypic differences can readily accumulate under strong selective pressures, bearing implications for efforts to prevent unwanted evolution in microbes.

## INTRODUCTION

Adaptation to strong selection pressures inevitably reduces genetic diversity. The fixation of adaptive alleles reduce variation not only at the locus where they occur, but also at all other genomic regions that are not separated from this locus by recombination^1^; a phenomenon referred to as a “selective sweep”^2,3^. Early models assumed that beneficial mutations were exceedingly rare, so selective sweeps would typically have a single origin and diversity would be drastically reduced (i.e., a “hard sweep”)^4,5^. Recent work, however, emphasizes that many natural populations can be large enough as to make beneficial mutations common, especially in microbes^6,7^. As a result, adapted populations often consist of multiple genotypes carrying alternative solutions to the same adaptive challenge (i.e., a “soft sweep”)^8,9^. The extent to which this genetic variation can have evolutionary consequences beyond the local selective conditions is crucial to many fundamental and applied problems in ecology and evolution. Examples include understanding biodiversity patterns along environmental gradients^10,11^, the tempo and mode of divergence and incipient speciation^12,13^, and the capacity of populations to adapt to new challenges^14,15^, crucial both for conservation efforts and for pest and pathogen control^16^.

A scenario in which this issue is particularly relevant is the fight against antibiotic resistance evolution, a major threat to public health worldwide^17^. Next-generation sequencing has revealed that bacterial populations adapted to antibiotics, both in experimental and clinical settings, are often comprised by multiple clones with different resistance determinants^6,18–20^. While presumably equivalent in fitness under the primary selective conditions, these different alternatives may display distinct liabilities and strengths across other clinically relevant conditions. This possibility has direct relevance to approaches that seek to apply thinking to prevent or eliminate antibiotic-resistant variants^21–24^. One such approach is prioritizing antibiotics for which resistance comes at a substantial fitness cost, either by reducing growth rates, transmission, or virulence, or by increasing sensitivity to the immune system or the host’s internal environment^25^. Another, more recently advocated strategy is the use of drug combination or cycling regimes that exploit trade-offs among antibiotics (i.e., collateral sensitivity), based on the common observation that resistance mutations to a given drug sometimes induce hypersusceptibility to other drugs^26,27^.

While promising, the success of these strategies in clinical practice hinges on the assumption that evolutionary outcomes are largely predictable; an assumption that cannot be taken for granted, especially under the variable conditions of real-world applications^28–30^. In fact, our current understanding of evolutionary genetics raises at least two important cautionary points to be considered. First, as many authors have already pointed out, resistance mutations are rarely unique or functionally identical; instead, they are better depicted as a pool of different classes with varying frequencies and phenotypic profiles^28,31–33^. As a consequence, the behavior of a “typical” mutant may not represent the entire distribution it comes from. Moreover, changes in population genetic parameters can alter which subset of mutations can be deemed as “typical”: experiments show that variations in population size, mutation bias and selection strength can result in different subsets of mutations becoming preferentially substituted during adaptation^6,28,34–37^.

A second, less appreciated cautionary point concerns epistasis (i.e., non-additive interactions among mutations). Epistasis is commonly observed among beneficial mutations, reflecting the many non-linearities in the mapping from genotype to fitness that can arise at structural, metabolic, and regulatory levels^38–41^. Of particular evolutionary relevance is strong epistasis, which can alter not only the magnitude but also the direction of fitness effects when mutations are combined (i.e., sign epistasis)^42^. Strong epistasis may pose two challenges to attempts to exploit trade-offs for applied purposes. First, since new mutations may interact idiosyncratically with previous ones, there is no guarantee that any trade-offs observed in a “typical” mutant will persist further along an adaptive pathway, making snapshot measurements of collateral sensitivities unreliable^28^. Second, since strong epistasis creates incompatibilities among entire sets of mutations, mutational paths become contingent on which mutations happen to occur first^43^. As a consequence, strong epistasis may lead to rapid divergence across multiple traits of lineages even within the same population, thus decreasing the likelihood that a “typical” mutant accurately represents how populations behave in secondary environments.

To explore how rapidly epistasis can drive phenotypic divergence both among and within mutational paths, here we turned to the highly tractable and relevant case study of enzyme-mediated antibiotic resistance. Antibiotic resistance enzymes are ideal model systems for several reasons. First, they are major actors underpinning resistance in clinical settings, posing significant concern due to their potential to spread rapidly across bacterial species^44,45^. Second, strong epistasis is typically pervasive among beneficial mutations within enzymes, as different residues often interact with each other either through direct physical contact or indirectly via altering stability and folding – indeed, much of our current understanding of the causes and consequences of intramolecular epistasis come from work done in the β-lactamase enzyme TEM-1^41–43,46–50^. Third, resistance enzymes often show substrate promiscuity across multiple antibiotics within the same class, and the evolution of increased activity against non-preferred substrates is a worrisome and well documented phenomenom in clinical settings^51,52^. As a result, these enzymes afford great opportunity to explore how adaptation to non-preferred substrates can proceed via alternative paths with distinct phenotypic profiles. Finally, for the most relevant enzymes, extensive knowledge exists on the structure-function relationship of many prevalent variants, aiding with the interpretation of laboratory evolution experiments.

Here we used directed evolution to assess the extent to which epistasis can drive the rapid divergence across multiple traits of the clinically relevant and globally distributed carbapenemase KPC-2 (*Klebsiella pneumoniae* carbapenemase-2)^53^. This class-A serine β-lactamase was originally named for its ability to hydrolyze carbapenems, last-resort drugs for treating multi-drug-resistant infections^54^. However, KPC-2 and its variants are also active against a wide range of other β-lactam compounds, including penicillins, monobactams, and cephalosporins^55,56^. This broad-spectrum phenotype, along with its rapid worldwide dissemination and high prevalence in Gram-negative bacteria, has established KPC-carbapenemases as a top priority for epidemiological surveillance in clinical settings^53,57^. KPC-2 is also noted for its versatility and high evolutionary potential^58^; with over 200 allelic variants reported in less than three decades since it was first described (bldb.eu^59^). These variants have mostly diverged through point mutations, although examples of small insertions and deletions also exist, and for many of the variants these changes are known to result in substantial alterations of their activity spectrum^58,60^.

One of the few yet relevant compounds for which KPC-2 shows poor activity is ceftazidime (CAZ)^61^, an important β-lactam cephalosporin commonly used in hospital and community settings to treat Gram-negative and some Gram-positive bacteria^62^. Leveraging this weakness, standard practice against KPC-bearing pathogens involves using CAZ in combination with β-lactamase inhibitors such as avibactam (AVI)^63–65^. However, despite the fact that CAZ-AVI combination has been introduced only recently (in 2015 in the USA and in 2016 in Europe), numerous reports have documented the emergence of resistant mutants after treatment, typically carrying mutations that alter the conformation of active-site loops to increase activity against CAZ^58,60,66,67^. Inspired by this natural case study, here we sought to characterize, via *in vitro* directed evolution, the diversity of mutational paths that enable KPC-2 to confer increased resistance to CAZ; placing especial emphasis on assessing how readily these paths can diverge not only in terms of genotypes, but also at multiple phenotypic levels.

## RESULTS

### Initial CAZ-resistance proceeds via mutations often seen in clinical variants

To explore the potential of KPC-2 to evolve high activity against CAZ, we began by cloning a wild-type *bla_KPC-2_* gene into a high-copy-number plasmid in *E. coli* and then used error-prone PCR to create 11 libraries with ∼10^3^–10^4^ randomly generated mutants (Methods). After overnight growth, we screened for CAZ-resistance phenotypes across a gradient of doubling concentrations of CAZ, starting from the Minimum Inhibitory Concentration (MIC) measured for the wild type (16 µg/ml) (Methods). From each library, we selected two colonies from the highest concentration of CAZ that sustained growth, which in this first round amounted to a 16-32 fold increase over the MIC of the ancestor (256-512 µg/ml). Sanger sequencing of these initial 22 colonies revealed a total of 38 single nucleotide polymorphisms in the *bla_KPC-2_* gene (1.54 ± 0.66 per isolate, mean ± sd), of which 15 were unique (Table S1). These nucleotide mutations translated into 34 amino acid substitutions (13 unique), arranged into nine different genotypes (Table S1, Figure 1A).

**Figure 1.**
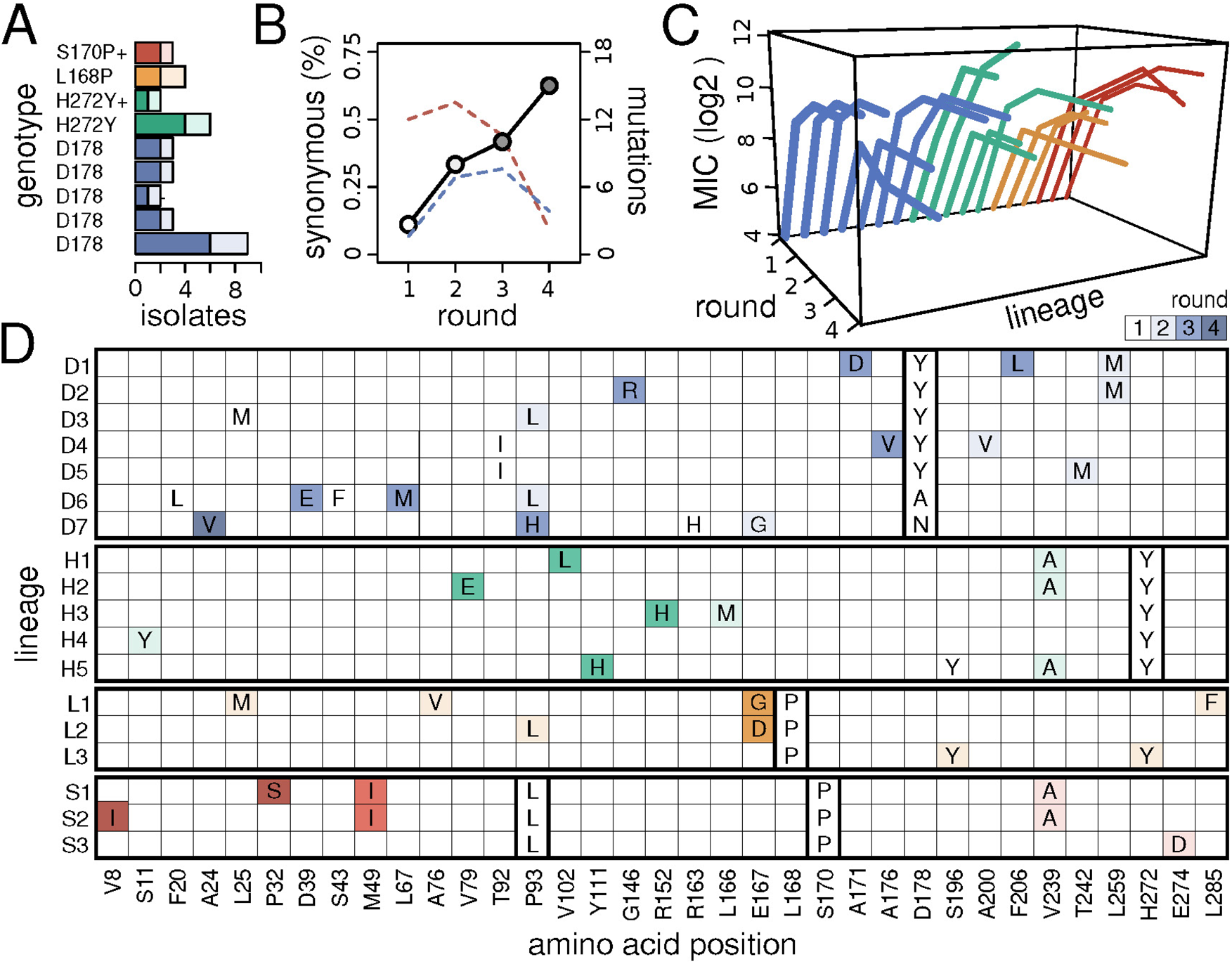
Mutational pathways recovered during KPC-2 directed evolution for increased CAZ resistance. **(A)** Abundance of the nine unique genotypes retrieved among the initial 22 colonies isolated in Round 1. Colors indicate grouping into four categories, according to mutations in positions previously associated with CAZ resistance. The presumed driver mutation is labeled on the y-axis, with a ‘+’ sign indicating additional accessory mutations (see Table S1). Darker shades represent total abundance across replicates, while lighter shades indicate independent occurrences. (B) Proportion of synonymous mutations across rounds (solid black line with circles; darker gray shades represent later rounds). The secondary y-axis displays the total number of synonymous (blue dashed line) and non-synonymous mutations (red dashed line). (C) Diversity of CAZ resistance trajectories across rounds and lineages, grouped and colored as in (A). (D) Horizontal alignment of the 18 lineages studied, highlighting only positions with mutations. Lineages are arranged, grouped, and colored as in (A), with darker shades representing later rounds. Presumed driver mutations are highlighted with thick-lined squares to underscore they are the basis for defining each group.

We next sought to classify the first-round mutants according to the mutations most likely driving the MIC increases. Clinical CAZ-resistant variants typically harbor mutations in three mutational hotspots^58,61^: the Ω-loop, forming the catalytic pocket floor; loop 240, forming the active site’s right wall; and loop 270, which undergoes conformational changes upon CAZ binding. Each of our initial 22 mutants carried mutations mapping to either the Ω-loop or loop 270. Consequently, we were able to define four classes of mutants based on positions known to feature CAZ-resistance mutations. The first class consists of mutants at position D178, located in the Ω-loop. This position was the most commonly mutated among the initial 22 colonies (13/22, 59%, Table S1), and was present in more than half of the initial genotypes (5/9, Figure 1A). This prevalence may be linked to being the only position in which a single mutation confers the highest observed MIC increase (D178Y, 32-fold), although mutational biases may also play a role^68^. Interestingly, reports show that this position is also the one most commonly harboring single point mutations among CAZ-resistant clinical variants^58^. We also observed substitutions at other positions within the Ω-loop, specifically L168 and S170 (2/22, 9% each, Table S1), and we took them as conforming different classes with potentially different evolutionary prospects. The final class comprises position H272, located in loop 270, the second most commonly altered among our mutants (5/22, 23%, Table S1). Of note, a mutant carrying substitution H272Y alone is known as KPC-3 in the literature^69^, the most prevalent clinical variant overall after KPC-2^70^.

### Alternative mutational paths drive KPC-2 evolution towards CAZ-resistance

Despite starting with 22 potential lines, the first round of directed evolution produced only nine different genotypes, which fell into just four groups based on likely CAZ-resistance mutations. Would these variants lead to markedly different evolutionary outcomes? To increase the chances of exploring less common mutational paths, we conducted a second round of directed evolution not only with the initial nine genotypes but also with six duplicates chosen from across the four groups. In particular, we added one line from each of the D178 and S170 groups (lines D5, S3, Table S2), and two lines from each of the H272Y and L168 groups (H3, H4, L2 and L3, Table S2). These latter groups were favored because they showed the smallest increases in MIC (16-fold), potentially a hallmark of suboptimal paths^50^. We subjected the resulting 15 lines to a new round of error-prone mutagenesis and screening, following the same conditions as in round one. The results contrasted to those seen before: while in round one we observed large and relatively homogeneous changes in MIC (16- and 32-fold increase), values in round two were much smaller and heterogeneous (from 0.5- to 8-fold increase, in doubling increments) (Table S2). This heterogeneity suggests that the lines have already begun to diverge substantially, with most still undergoing adaptation to different extents (10/15), a few reaching a plateau (4/15, lines D5, H3, H4 and L1, Table S2), and one line even experiencing a MIC decline, likely due to the random sampling of a mutant with a slightly deleterious mutation (A200V, line D4, Table S2).

To guide our selection for the third round of directed evolution, we considered three criteria. First, we included the 10 lines that kept sustaining adaptation thus far. Second, we duplicated two lines with the highest MIC values to explore potential alternative routes to maximum CAZ resistance (D1 and H1, with MICs of 1024 and 2048 µg/ml, a 64- and 128-fold increase over the wild-type, respectively). Third, we included three out of the four lines that had reached a plateau in the previous round, as well as the line that declined in MIC. The reason for continuing these lines was to investigate what drove their apparent halt in adaptation: was it due to MIC-increasing mutations being just infrequent, or rather because they were essentially unavailable? After these considerations, we proceeded with a total of 16 lines for a third round of error-prone mutagenesis and screening, under the same conditions as before. In stark contrast to previous rounds, here the majority of lines showed no further adaptation (14 out of 16), and the only two lines that did adapt exhibited only a modest 2-fold increase in MIC (H2 and S1, Table S2).

Despite MIC increases mostly coming to a halt overall after round three, we decided to subject a fraction of the lines to a final, fourth round of directed evolution. An obvious choice for this last round were the two lines that were still adapting, even if modestly. These lines were duplicated to further explore potential routes to maximum CAZ resistance, although for H2 no further mutations appeared so the duplicate is not included in Table S2. We also included five lines across groups that, despite having already reached a plateau, displayed either some of the highest (D6, D7 and H5, Table S2) or the lowest MICs increments overall (D4 and L1, Table S2). We included these lines, representing plateaus of contrasting heights, to explore whether the non-synonymous mutations acquired in the previous round could act as cryptic stepping-stones towards the acquisition of further adaptive mutations^71^. In total, we subjected nine lines to a fourth round of error-prone mutagenesis and screening. None of the lines presented any further increase in MIC (Table S2).

Figure 1 summarizes the basic features and results of the four-round, directed evolution experiment. Three major takeaways are possible. First, at least four distinct mutation sites can mediate large-effect, first-step resistance mutations towards CAZ resistance (Figure 1A). These sites, however, show a markedly uneven representation, with one site accounting for 59% of the first-step mutants (D178), presumably because it enables the highest single-step MIC increase. Second, while the proportion of synonymous mutations showed no evidence of plateauing (Figure 1B), the MIC trajectories slowed down almost in parallel across groups, with most lines reaching a plateau by round three (Figure 1C). This contrast implies an excess of non-synonymous mutations during the initial versus the later stages of plateaus, potentially indicating that plateaus are not adaptively stagnant. Could it be that these non-synonymous mutations, invisible in terms of MIC levels, are mediating the cryptic adaptive evolution of other fitness-related traits?

The third takeaway is that first-step mutations directed adaptation through distinct, mostly non-overlapping sets of adaptive mutations, indicating that strong epistatic incompatibilities are pervasive^43^. Indeed, only 6 out of the 35 sites featuring mutations were hit in more than one group (17%, L25, P93, E167, S196, V239, and H272; Figure 1D), with only one site hit in three groups (P93), and none in all. However, this stark genetic divergence across groups translates into just moderate levels of divergence in CAZ-resistance (Figure 1C). Indeed, only groups L168 and S170 show statistically significant differences in endpoint MIC levels (*P = 0.024*, Student’s t-test on log2 transformed values). In other words, the different groups may be exploiting alternative solutions at the molecular level, and yet these solutions converge to similar values of the major fitness-related trait in the selective conditions. While this convergence could be interpreted in terms of global physio-chemical limits to enzymatic activity, it is still possible that the different genotypes behave differently across secondary traits that matter to fitness in other environments. Could the high degree of genetic divergence translate into markedly different properties beyond the original selective conditions?

### Collateral sensitivities rapidly diverge during CAZ-resistance evolution

We next sought to assess the extent to which the different lines adapted to CAZ may show cross-resistance or cross-sensitivity to other beta-lactam antibiotics, as cycling among beta-lactams has been proposed as a convenient intervention to curb resistance evolution^72–74^. To this end, we chose three compounds commonly used to treat Gram-negative infections, with different degrees of chemical similarity to CAZ^75^. These include cefotaxime (CTX), which belongs to the same subclass as CAZ (3^rd^ generation cephalosporins); cefoxitin (FOX), a 2^nd^ generation cephalosporin; and imipenem (IMI), an important representative of the carbapenem class. Figure 2A reveals a general trend for CAZ-adapted lines to become more susceptible than the ancestor to the other antibiotics, although the fraction of lines showing this increased susceptibility varies considerably (92.9% for IMI, 78.6% for FOX, and 57.1% for CTX). Another relevant point is that MIC trajectories for the other antibiotics show less clear plateaus than those seen in CAZ, with frequent fluctuations occurring at rounds where CAZ MIC values have already stabilized. This observation provides further evidence that the non-synonymous mutations detected in plateaus have some functional relevance: these changes are probably driving the adaptive evolution of other fitness components and, while not affecting CAZ MIC values, manifest in the MIC values for other drugs.

**Figure 2.**
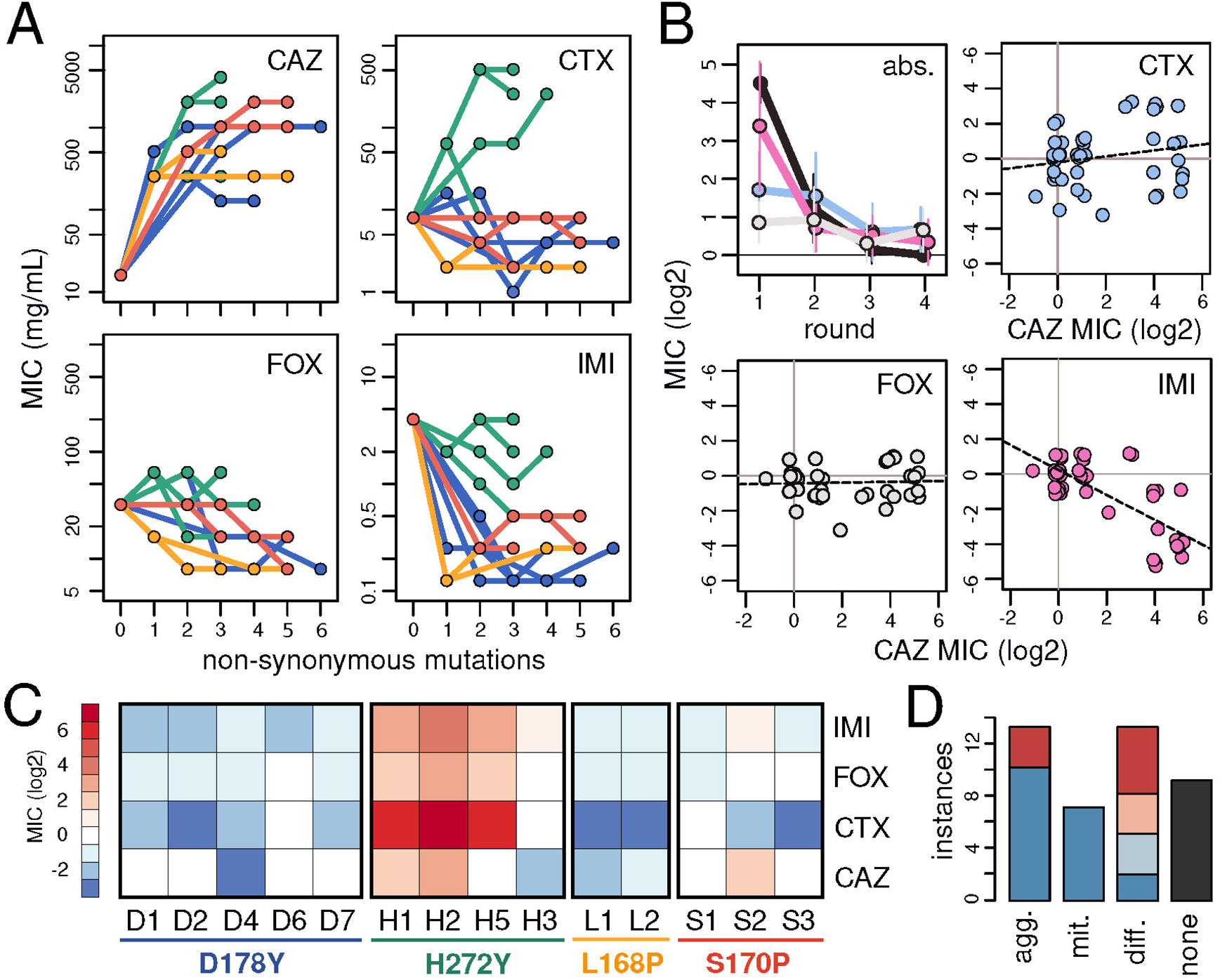
Collateral sensitivity toward other β-lactams rapidly diverges during CAZ-resistance evolution. (A) Resistance trajectories against the number of non-synonymous mutations accumulated across rounds for each lineage, with colors indicating the grouping based on positions associated with CAZ resistance: blue (D178Y), green (H272Y), yellow (L168P), and red (S170P). Panels show MIC measurements for CAZ, cefotaxime (CTX), cefoxitin (FOX), and imipenem (IMI), as labeled in the upper right corner. (B) Patterns of resistance across rounds. The first panel shows the average overall change in magnitude for each β-lactam, with error bars representing standard deviation. Colors denote averages for CAZ (black), CTX (cyan), FOX (grey) and IMI (pink). The remaining panels show correlations between individual MIC changes for CAZ and the other antibiotics, with dashed lines indicating the best fit from linear regression. (C) Variation in collateral sensitivity among and within lineages. Lineage labels are color-coded as in (A). Resistance values for endpoint isolates are represented following a color gradient (blue, increased sensitivity; red, cross-resistance). Note values denote relative variation compared to all lineages. (D) Divergence in collateral sensitivity along mutational pathways. Values indicate the number of lineages for which the full path has aggravating, mitigating, different, or no effects on the first-step mutant resistance. Blue indicates that the first step caused increased sensitivity; red indicates it caused resistance. Shaded colors represent cases where the endpoint profile is neutral compared to the wild-type.

To further investigate the observed tendency towards collateral sensitivity, we examined how the overall magnitude of collateral effects changed through rounds. Since most high-impact CAZ resistance mutations were incorporated early on, it is reasonable to expect the most pronounced collateral effects to occur early as well^76,77^. Figure 2B confirms this expectation, but also shows that while these effects tend to decay over time (right top panel), they do at varying rates. Indeed, there is a significant negative correlation for strong collateral effects occurring mostly at the beginning of the experiments for CTX and IMI (Pearson’s *R = −0.68* and *-0.42*, respectively; *P < 0.01* for both), but not for FOX (*R = −0.27*; *P = 0.068*). We then focused on collateral effects at the level of each individual mutational step, looking for correlations between the change in MIC for CAZ versus the other antibiotics. The remaining panels in Figure 2B reveal that only individual collateral effects in IMI exhibit a significant correspondence with the primary effects seen in CAZ (*R = −0.74*; *P < 10^-7^*), whereas the correspondence for CTX and especially FOX is virtually absent (*R = 0.21* and *R = 0.05*; *P = 0.180* and *0.741,* respectively). The discrepancy in statistical significance observed for CAZ between absolute, global effects and actual, individual ones likely reflects the substantial degree of divergence that lines experienced in this phenotype, with some developing cross-sensitivity and others cross-resistance (Figure 2A).

We next sought to assess the extent to which epistasis can cause heterogeneity in collateral sensitivity profiles. This heterogeneity can manifest in two dimensions: among lines, and over time along each line. First, we focused on heterogeneities among endpoint isolates, examining differences both among and within each group. Figure 2C shows that endpoint sensitivity profiles vary markedly among groups, primarily due to the tendency of lines in the H272 group to exhibit MIC values that consistently rank highest across antibiotics (P < 0.002, two-sample Wilcoxon test on ranked data). However, we note that a few lines deviate from the typical profile of their group (H3 and, to a lesser extent, S2). This deviation illustrates that, even when initial mutations markedly determine the properties of subsequent adaptive steps, some secondary mutations can cause pathways to branch away from the overall profile of the main route. Interestingly, in both H3 and S2, these deviations involve mutations that do not affect CAZ MIC values (Table S2), illustrating how cryptic changes during plateaus can have unexpected consequences in phenotypic profiles.

Finally, and in line with the last point, we sought to quantify how much divergence typically happens over time along a single mutational path. To this end, for each line and secondary antibiotic combination, we classified first-step mutants as cross-resistant, neutral, or cross-sensitive compared to the ancestor; and then assessed how this classification changed for endpoint mutants. Figure 2D shows the extent and direction to which endpoint isolates deviate from the collateral sensitivity profile of the first-step mutant of their respective lines. We found that the most common paths were those along which sensitivities changed classification (33%, 14/42), followed by paths along which effects were aggravated (31%, 13/42), paths with no change (19%, 8/42), and paths along which effects were mitigated (16.7%, 7/42). Notably, all paths with mitigating effects involved cross-sensitive first-step mutants.

### Molecular traits rapidly diverge during CAZ-resistance evolution

Thus far, we have interpreted our results in terms of MICs, which are macroscopic readouts that arise from the aggregate contributions of multiple molecular-level traits. Among these, the traits typically considered to shape MIC the most are those directly affecting how efficiently an enzyme converts substrate into product (i.e., the “catalytic efficiency”, the ratio of the turnover number over the substrate affinity, *k_cat_/K_m_*)^49,78^. However, recent work revealed a major role for the multiple traits that affect the different aspects of the beta-lactamase’s life cycle, including its exit from the ribosome as a precursor polypeptide, translocation across the inner membrane, folding once in the periplasm, and their eventual degradation by proteases^79–83^. The intuition for this importance is that, regardless of how efficient the beta-lactamase may be, these factors ultimately determine effective reaction rates by controlling the steady-state abundance of functional enzymes in their natural workplace^83^.

Although the specific molecular traits altered are often difficult to pinpoint, an aggregate readout of all these traits can be obtained by directly measuring enzyme abundance stability in the periplasm^79,81^. Indeed, these periplasmic stability measurements have successfully helped explain MIC differences among closely related enzyme variants^83^, and most revealingly, among the same enzyme across different hosts^82^. Moreover, beyond MICs, these traits largely control fitness costs in the absence of drugs, as alterations in any stage of the enzyme’s life cycle can interfere with other cellular processes, such as saturating the translation, membrane translocation, and folding machinery, or triggering stress responses due to the aggregation of misfolded proteins^84,85^. This impact on fitness in the absence of strong selective pressures for resistance is expected to be crucial for the long-term success of enzyme variants under ever-changing conditions^82–84^.

To gain insights into how epistasis may have driven divergence in molecular-level traits during CAZ adaptation, we estimated the catalytic efficiency (*k_cat_/K_m_)*, periplasmic abundance stability and growth rates of a fraction of the lines. The results are summarized in Figure 3. We first examined the catalytic efficiency of purified extracts for several variants (Methods), including all steps in the line that achieved the largest MIC increase (H2), an alternative endpoint isolate deviating from this line (H1), and three high MIC representative isolates from across the other groups (D1, S1 and L2). Figure 3A shows variations in catalytic efficiency spanning more than three orders of magnitude, including one line in which almist no increase relative to the ancestor was detected (D178Y). These observations contrast to the fact that CAZ MICs increased substantially in all lines, and that in endpoint isolates MIC values vary less than 10-fold (512 – 4096 µg/ml, Table S1). Similar results were obtained using hydrolysis rate estimates from a colorimetric assay, an alternative method for assessing pure enzymatic activity^60^ (Figure SX) (Methods). Finally, we ask if at least at the level of each individual step, the effects on MIC and catalytic efficiency show any degree of correlation (Figure 3A, bottom panel). In line with previous studies^49,81,83^, no statistically significant correlation was detected (*R = 0.01; P = 0.98*).

**Figure 3.**
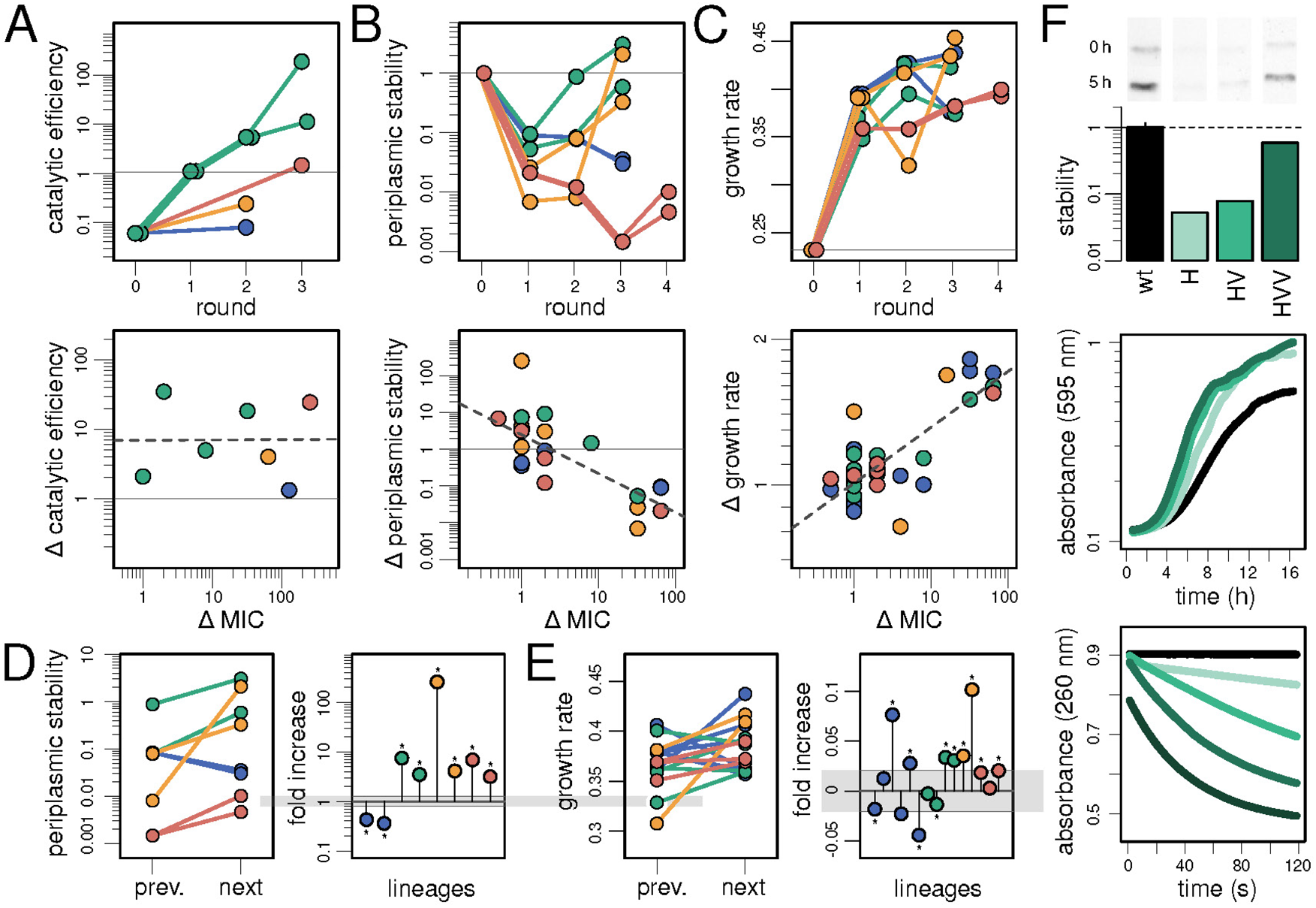
Molecular traits rapidly diverge during CAZ-resistance evolution. (A) Top panel: catalytic efficiency (*k_cat_/K_m_*) across rounds for selected enzyme variants, expressed as fold change relative to the ancestral KPC-2 enzyme. Colors indicate initial CAZ resistance mutations: blue (D178Y), green (H272Y), yellow (L168P), and red (S170P). Bottom panel: correlation between individual changes in *k_cat_/K_m_* and MIC, with best-fit (dashed line) and ancestral value (solid line). (B) Top panel: stability of enzyme abundance in the periplasm across rounds for the same variants as in Figure 2. Values represent Western blot intensity changes over five hours of growth, relative to ancestral KPC-2, expressed relative to the ancestral KPC-2 enzyme (see F). Color coding as above. Bottom panel: correlation between individual changes in periplasmic stability and MIC. Dashed line indicates the best fit from linear regression; horizontal solid line stability of the ancestral enzyme. (C) Top panel: growth rates in sub-MIC concentrations of CAZ across rounds for the same variants as in (B). Bottom panel: correlation between individual changes in growth rate and MIC. Dashed and solid lines show best fit and reference values as above. (D) Change in periplasmic stability between consecutive mutants within plateaus (left). Change expressed as fold increase (right); gray bar represents measurement uncertainty (mean ± 2 x standard deviation for the wild-type). Asterisks indicate statistical significance. (E) Similar analysis as in (D), but showing changes in growth rate between consecutive mutants within each plateau. (F) Top panel: example Western blot quantifying periplasmic stability, with a bar plot showing the log intensity ratio of initial (1 h) and final (5 h) bands for each variant. Middle panel: example growth rates in sub-MIC CAZ concentrations for the same variants. Bottom panel: example CAZ concentration decay from the colorimetric assay for the same variants.

We next examined the stability of enzyme abundance levels in the periplasm (i.e., in-cell kinetic stability, dubbed this way to differentiate it from standard, *in vitro* thermal stability^86^). To this end, we added a hemagglutinin tag to wild-type KPC-2 and several evolved variants, and used Western blot to quantify the change in their relative abundance in the periplasm over time (Figures SX and SY) (Methods). For this study, we chose two high-MIC lines from each group, including all the individual mutants from each line. Figure 3B reveals a common pattern in which periplasmic stability systematically drops during the first steps of adaptation, later recovering to different extents in all lines, except from the D178-derived ones. In addition, while here again the values for the endpoint isolates vary widely (four-orders of magnitude), analyses at the level of each individual mutation step show a clear negative correlation between the effects on MIC and on periplasmic stability (*R = −0.7*; *P = 0.08*) (Figure 3B, bottom panel). Interestingly, the largest individual effect on stability in this panel corresponds to a mutation that does not change MIC (L1, E167G, Table S1), reinforcing the idea that adaptive evolution did not stop once populations reached the MIC plateaus. Further support for this idea comes from the observation that endpoint isolates from S170P-derived lines exhibited an altered molecular weight at later incubation times (Figure SX). Proteomics confirmed the presence of the enzyme, suggesting the possibility of structural or post-translational modifications that may enhance stability by facilitating binding to other components of the periplasm.

We then measured growth rates as a proxy to capture any collateral fitness costs resulting from the evolution towards CAZ resistance^84,85^. We conducted the assays at sub-inhibitory concentrations of CAZ not only to be able to include the ancestor, but also to assess the severity of the potential costs under conditions where there is still a low, yet consequential, demand for the enzymes to perform their physiological function^87^. We first estimated growth rates for the same two high-MIC lines from each group used in the periplasmic stability study, including all individual mutants along lines. Figure 3C shows a general trend whereby growth rates become larger with each round of evolution, which is expected simply due the higher protective capacity of the evolved enzymes. However, endpoint growth rates exhibit variations that do not align with MIC difference. This is best exemplified by line L1, which displays the highest growth rate despite having the lowest MIC (256 µg/ml, Table S1). Similarly, S170-derived lines show the lowest growth rates even though they have some of the highest MIC values (1024 and 2048 µg/ml, S1 and S2, respectively, Table S1). At the level of each individual mutation step, we found a significant positive correlation between the effects on MIC and on growth rate (*R = 0.77*; *P = 0.02*) (Figure 3C, bottom panel). Lastly, it is worth noting line L2, with undergoes a large drop in growth rates in round three, followed by a full recovery in the next round. This recovery involves a mutation that does not alter MIC (E167D, Table S1), again pointing at the non-static nature of plateaus.

Finally, we wanted to characterize the patterns of change occurring specifically during the plateaus. We first examined periplasmic stability, looking at the direction of change between consecutive mutants within each plateau. Figure 3D shows that periplasmic stability increases during plateaus in all lines, except for the D178-derived ones (6/8, Wilcoxon Rank Sum test, P > 0.05 in all cases). We then turned our attention to growth rates, for which we extended growth curve measurements to all mutants in plateaus across lineages (Figure SZ). Figure 3D reveals that, despite MIC stasis, a majority of lines exhibit a significant increase in growth rate (8/15), while only three lines show significant decreases (Wilcoxon Rank Sum test, P > 0.05 in all cases). Overall, these results point at the idea that adaptive evolution continues across multiple molecular traits even after CAZ-resistance levels stabilize. Moreover, a glance at the location and predicted effects of mutations over rounds provides further support for this notion (Figure 4). Initial mutations involve disruptive substitutions near the active site, likely reshaping its physical and chemical properties to enhance enzymatic activity. Later mutations tend to occur farther away and have milder effects, possibly compensating for the stability loss and collateral fitness costs incurred by the initial activity-enhancing mutations. Similar patterns of mutations falling progressively away from the active center have been observed in directed evolution experiments with other proteins^88,89^.

**Figure 4.**
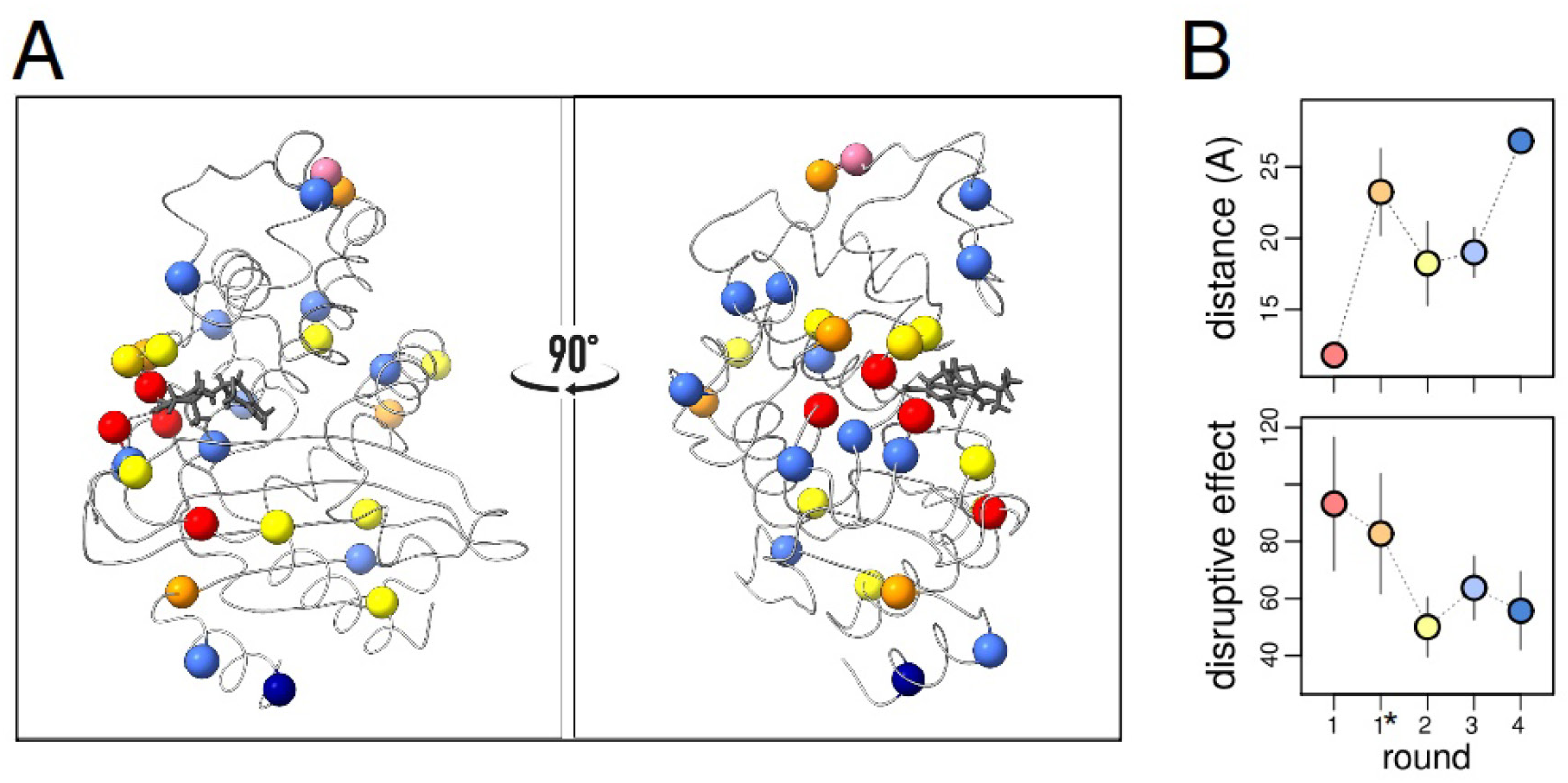
Position and properties of accumulated mutations across rounds. **(A)** Diagram of the KPC-2 β-lactamase backbone with CAZ bound at the active site (shown as sticks). Positions where mutations were observed are represented as spheres, colored according to the round in which they arose: red (Round 1, presumably driver mutations), orange (Round 1, accessory), yellow (Round 2), light blue (Round 3), dark blue (Round 4). The purple sphere marks the P94 substitution, present in all rounds of mutagenesis. **(B)** Average atomic distance of mutations from CAZ’s center of mass (top), and their Grantham score (physiochemical distance) as a proxy of their disruptive effect per round. Circle colors follow the same scheme as in (A).

## DISCUSSION

In this study, we sought to assess the extent to which strong epistasis can lead lineages to diverge not only at the genotypic level, but also across other traits that may be relevant in alternative conditions. To this end, we characterized the diversity of mutational paths through which the globally distributed KPC-2 carbapenemase can evolve increased activity towards a poor substrate, the beta-lactam antibiotic CAZ. Our results show that KPC-2 can readily adapt through different mutational paths with markedly distinct phenotype properties, including differences in catalytic efficiency, periplasmic stability, collateral fitness costs, and susceptibility to other antibiotics. The speed and degree of divergence observed, both among and within lines, are remarkable considering that we conducted the experiments under identical and well-controlled laboratory conditions; and bear implications for designing strategies to prevent antibiotic resistance, and for understanding natural evolution in complex environments.

Can we rationalize the molecular basis for why the initial CAZ-resistance mutations lead to such divergent outcomes? We classified the lines into four groups based on the positions altered in the initial CAZ-resistance mutations. These positions, however, vary in how frequently they are altered among all natural KPC-2 variants: at present, position H272 retrieves the most hits (47%) in the BetaLactamase DataBase (http://bldb.eu)^59^, followed by D178 (26%), L168 (15%), and S170 (9%). Reflecting this unequal prevalence, detailed studies on the molecular mechanisms of resistance are available only for positions D178 and H272. Substitutions N and Y at D178 increase Ω loop flexibility, facilitating the rapid entry of CAZ into the active site^90,91^. However, this flexibility reduces hydrolysis rate due to the displacement of essential hydrolytic residues. It is not surprising, therefore, the reductions in both stability and hydrolitic efficiency that we observed (Figure 3A-B, Figure SX). Interestingly, increased resistance in D178 mutants is attributed to a mechanism referred to as a “kinetic trap”, whereby the swift entry of CAZ into the reaction pocket reduces to safe levels the amount of free molecules available for inhibiting penicillin-binding proteins^91^. In stark contrast, substitution Y at H272 enhances hydrogen bonding with CAZ’s oxyimino side chain, directly increasing the hydrolysis rate^60^. Additionally, H272Y is thought to aid CAZ entry into the active site without altering the Ω loop’s flexibility, which is achieved by the repositioning of the surrounding 240 and 270 loops^92^. So to sum up, these well-studied positions represent fundamentally different mechanisms for enhancing beta-lactamase activity; being therefore reasonable to expect that they may give rise to distinct phenotypic profiles.

Among the implications of our findings, the most immediate pertains to current efforts to exploit collateral sensitivities in the fight against antibiotic resistance^93^. Our work demonstrates that widespread strong epistasis can drive the rapid and substantial divergence of resistance profiles across different antibiotics, even in parallel replicates evolving under the same conditions. Moreover, this divergence can occur not only among lines initiated from distinct mutations, but also along the sequential steps that comprise each line. These observations reveal extra challenges to the endeavor of anticipating evolutionary responses to antibiotics^28–33,35^. For instance, resistance trade-offs may markedly differ among seemingly similar populations, due to being dominated by the descendants of different mutants that happened to be the first to occur. Furthermore, heterogeneities in resistance are also expected among individuals within a single population, provided its size is sufficiently large. To further complicate matters, the observed trade-offs may not even be stable over subsequent steps of adaptive evolution. These phenomena call into question the reliability of snapshot characterizations of “typical” mutants as indicators of populations’ responses to future treatments^28^. Thus, this work adds to the cautionary evidence suggesting that collateral sensitivity strategies, while promising, need to be carefully tailored to take into account the many factors influencing evolutionary repeatability^28–33,35^. This body of observations calls for a careful and thorough characterization of the spectrum of resistance mutations, within the limits of technical feasibility, if we are to ensure the best success of these strategies.

Our findings also bear implications for understanding the natural evolution of antibiotic resistance enzymes, as well as the distribution patterns of variants across regions and species. Bacterial populations, especially in clinical pathogens, are regularly exposed to multiple antibiotics with concentrations that can change unpredictably over space and time^87,94–96^. Since phenotypic profiles are strongly determined by the chance occurrence of initial mutations, which variants will prevail in a particular setting will be heavily contingent upon the specific order and intensity of the antibiotic pressures encountered by the population. Small population sizes or transmission bottlenecks will further amplify this historical contingency, simply due to the stochastic sampling of different variants with potentially distinct properties in other environments^28,50^.

Moreover, besides differences in antibiotic resistance profiles, we have shown that strong epistasis can readily lead to variants with substantial differences in periplasmic abundance and collateral fitness costs, traits that presumably reflect how enzymes interact with various cellular components throughout their life cycle^82–85^. These differences are expected to play a role in the scenario described above, favoring the selection of the least costly variants when antibiotics are at very low concentrations or even absent^84,85^. On the long run, these differences are probably critical for the evolutionary success of variants during their journey across multiple hosts^82,83,97^. Indeed, while we have presented evidence that variants interact differently with the cellular components of just one host, these differences may be amplified or mitigated in other hosts with their own idiosyncrasies in their cellular components. Thus, strong epistasis might fuel the rapid association of certain variants with specific hosts, a particularly intriguing possibility given how important horizontal gene transfer is in the dissemination of antibiotic resistance genes^44,45^.

Lastly, it is pertinent to ask whether the strong degree of epistasis-driven phenotypic divergence observed here with KPC-2 is peculiar to this enzyme, or if, conversely, similar patterns can be expected in other model systems. KPC-2 has been previously hypothesized to be more evolvable than other class A beta-lactamases because its thermal stability is significantly higher than other model enzymes like TEM and CTX-M (66.5°C versus 51.5°C and 51°C, respectively)^60^. High thermal stability enables enzymes to accumulate destabilizing beneficial mutations while maintaining proper folding and functionality, a property shown to boost evolvability in experiments with different systems^98,99^. Thus, compared to other class A beta-lactamases, an intriguing and testable possibility is that KPC-2’s enhanced stability may facilitate the acquisition of first-step mutations with a broader and more diverse range of molecular mechanisms for increasing enzymatic activity, being thus more prone to lead to divergent outcomes across multiple traits.

Beyond single proteins, it remains to be determined whether epistasis-driven phenotypic divergence is a general phenomenon in bacterial rapid adaptation to novel conditions. For instance, strong epistasis is common in the well-known Long Term Evolution Experiment with *E. coli*, in which 12 populations have been adapting to a low-glucose environment for over 75,000 generations^100^. However, the level of divergence across secondary environments that lineages experienced is relatively moderate, and it typically took thousands of generations to become manifest^101,102^. One possible explanation for this difference in breadth and pace is that in the LTEE selection pressures are milder. In addition, the functional target size for adaptation in the LTEE involves multiple genes, and therefore adaptation is not entirely dominated by the biophysical details of a single protein. Supporting this interpretation, a similar experiment that adapted 114 lines of *E. coli* to a nearly lethal temperature found much higher levels of divergence within just 2000 generations^103^. In this study, lines diverged along two epistatically incompatible mutational paths involving strongly adaptive mutations in key hubs of gene expression regulation. Notably, one path led to collateral resistance against rifampicin, an antibiotic never present in the medium^104^, providing a clear illustration of epistasis-driven divergence during genome-wide evolution. Further research is needed to clarify the factors that determine the degree to which strong epistasis can produce markedly different outcomes across environments.

## METHODS

### Bacterial strains and plasmids

*E. coli* TOP10 strain (Invitrogen, Massachusetts, USA) and the pUC-derived, high-copy number pCR-blunt II-TOPO vector (Invitrogen), carrying a kanamycin resitance gene, were used in the random mutagenesis experiments, susceptibility testing, growth curves and periplasm extract collection. *E. coli* Origami 2(DE3) strain (Novagen, Fontenay-sous-Bois, France) and pET-28a plasmid (Novogen) were used for protein expression and purification.

### Random mutagenesis

Wild-type *bla_KPC-2_* gene was amplified from a reference strain obtained from Dr. Naas laboratory^105^ (env-KPC-2_Fw: 5’ TTC AAA CAA GGA ATA TCG TTG ATG TCA CTG TAT CGC CGT CT 3’, env-KPC-2_Rv: 5’ AAT AGA TGA TTT TCA GAG CCT TAC TGC CCG TTG ACG CCC A 3’), cloned into the pCR-blunt II-TOPO vector (Invitrogen), and expressed in *E. coli* TOP10 (Invitrogen). Random mutagenesis was performed using the GeneMorph II EZClone Domain Mutagenesis Kit (Agilent) according to the manufacturer’s instructions. The *bla_KPC-2_* gene was amplified with a similar pair of primers as above but sparing the start of the open reading frame (PRE-KPC-2_Fw: 5’ TTC AAA CAA GGA ATA TCG TTG 3’, PRE-KPC-2_Rv: 5’ AAT AGA TGA TTT TCA GAG CC 3’) and using Mutazyme II DNA polymerase, which according to the manufacturer, generates a low-bias mutation spectrum with similar mutation rates at A’s/T’s and G’s/C’s. To achieve a low mutation rate (0–4.5 mutations/kb), 500 ng of target DNA and 30 PCR cycles were used. The resulting variant library was purified using the GeneJET PCR Purification Kit (ThermoFisher). A second PCR, using the mutant library as megaprimers and the pCR-blunt II-TOPO (bla_KPC-2_) plasmid as the template, was performed. In the next step, the template was digested with Dpn1 (ThermoFisher), and the PCR product was transformed into *E. coli* TOP10. Transformants were incubated overnight at 37°C in LB broth supplemented with 50 µg/ml kanamycin to ensure plasmid retention. Mutant selection was performed in Mueller Hinton (MH) broth (ThermoFisher). Four tubes containing 20 mL of MH broth with increasing concentrations of ceftazidime (CAZ) (ThermoFisher) were used. The first concentration corresponded to the midpoint of the minimal inhibitory concentration (MIC) conferred by KPC-2, and subsequent concentrations were doubled. Inocula were adjusted to 5 × 10⁵ CFU/mL, and cultures were incubated overnight at 37°C. The tube with the highest CAZ concentration showing visible growth was plated on LB agar with 50 µg/mL kanamycin. For the first mutagenesis round, two randomly selected colonies were sent for Sanger sequencing using M13 primers (M13_Fw 5’ GTA AAA CGA CGG CCA G 3’, M13_Rv 5’ CAG GAA ACA GCT ATG AC 3’), which were also used in subsequent rounds.

### Susceptibility testing

MIC values for *E. coli* TOP10 strains carrying KPC mutants were determined using the broth microdilution method following EUCAST guidelines. In brief, 96-well plates were used with a final volume of 200 µL per well. In the first well, 100 µL of MH broth and antibiotic stock (final concentration < 25% of 100 µL) were added to achieve the highest concentration tested. Serial two-fold dilutions of the antibiotic were then prepared. Finally, 100 µL of inoculum (1 × 10⁶ CFU/mL) from an overnight culture was added. MIC values were assessed after 18 hours of incubation at 37°C.

### Growth Curves

A 300 µL volume of LB broth containing 8 µg/mL ceftazidime was inoculated with *E. coli* TOP10 expressing KPC-2 and its variants and placed in 100-well honeycomb plates. Incubation was carried out at 37°C for 20 hours using the Bioscreen C Pro Microbiological Growth Analyser (Labsystems, Helsinki, Finland), with OD600 values measured every 10 minutes. Five replicates were used per variant. Growth rate values were determined from the maximum slope of the natural logarithm of optical densities over time.

### Protein purification

The *bla_KPC-2_* gene and variants’ fragments corresponding to the mature β-lactamase were cloned into the expression vector pET28a, using the PCR generated fragment with primers NdeI-KPC-2-sps_Fw (5’ CAT ATG CTG ACC AAC CTC GTC GCG GA 3’), BlpI-KPC-2_Rv (5’ GCT CAG CTT ACT GCC CGT TGA CGC CCA AT 3’) and the NEBuilder® HiFi DNA Assembly Cloning Kit (New England BioLabs®Inc, United Kingdom), following the manufacturer’s instructions. Recombinant plasmids were transformed into chemocompetent *E. coli* Origami 2(DE3) and the transformant were selected in LB agar plates supplemented with 50 µg/ml kanamycin. An O/N culture of *E. coli* Origami 2(DE3) harboring pET28a(*bla_KPC-2_* gene or *bla_KPC_*-mutant genes) was used to inoculate 2 L of LB broth containing 50 µg/L kanamycin. Bacteria were cultured at 37°C until reaching an OD of 0.6 at 600 nm. Expression of KPC-2 and variants was induced O/N at 25°C with 0.1 mM IPTG. The *bla_KPC-2_* gene and its mutants were purified in one step pseudo-affinity chromatography using a NTA-Nickel column (Cytiva, Freiburg, Germany). Protein purity was estimated by SDS–PAGE, pure fractions were pooled and dialyzed against 20mM Hepes SO4K2 50 mM buffer (pH=7) and concentrated by using Vivaspin® columns (Cytiva, Freiburg, Germany). Protein concentration was determined by Bradford Protein assay (Bio-Rad, Marnes-La-Coquette, France).

### Steady state parameters

Kinetic parameters of purified KPC-2 and its mutants were determined at 25°C in 100 mM sodium phosphate buffer (pH=7). The *k_cat_*and *K_m_* values were determined by analyzing hydrolysis of β-lactams under initial-rate conditions with an ULTROSPEC 2000 model UV spectrophotometer (GE Healthcare). Kinetic parameters were calculated using the non-linear regression of the Michaelis–Menten equation *v = kcat[S]/(Km+[S]).* The β-lactams were purchased from Sigma–Aldrich (Saint-Quentin-Fallavier, France). The velocity of ceftazidime hydrolysis could not be saturated by measurable concentrations due to a high *K_m_*. Thus, the second order rate constant at steady-state, *k_cat_*/*K_m_*, was determined by fitting the progress curves to the equation *v = k_cat_/K_m_[E][S]*, where *[S] << Km*^60^.

### Periplasmic stability

A C-terminal hemagglutinin tag was added to the proteins expressed by *E. coli* TOP10 strains harboring pCR-blunt II-TOPO (*bla*_KPC-2_ gene and variants) by using PRE-KPC-2_Fw and KPC-HA-STOP_Rv primers (5’ TTC AAA CAA GGA ATA TCG TTG 3’, 5’ TTA TGC ATA ATC CGG AAC ATC ATA CGG ATA CTG CCC GTT GAC GCC C 3’). Extraction of periplasmic proteins was adapted from^86^. *E. coli* TOP10 pCR-blunt II-TOPO strains (*bla*_KPC-2_ gene and variants) were incubated in LB broth supplemented with 50 µg/L kanamycin until reaching a OD_600_ value of 0.30, which was taken as T_0_. Two ml aliquots of the cultures were then taken at 1, 2, 5 and 10 h from T_0_. The aliquots were pelleted, washed with 20 mM Tris-HCl (pH 8.5) and 150 mM NaCl. To normalize the number of cells, the volume according to *V= (2 ml x 0.064 OD_600_)/OD_600_* was resuspended in 1ml of 20 mM Tris-HCl (pH 8.5), 0.1 mM EDTA, 20% w/v sucrose, 1 mg/ml lysozyme (Sigma-Aldrich, protein ≥90%), 0.5 mM phenylmethylsulfonyl fluoride (PMSF), incubated with gentle agitation at 4 °C for 30 min. After centrifugation at 15000 g for 2 min at 4°C, the supernatant containing the periplasmic fraction was collected and store at 4°C. Levels of C-terminal HA tagged proteins were determined in the periplasmic fraction, by SDS–PAGE, transferred to iBlot™ 3 Transfer Stacks, nitrocellulose (ThermoFisher), followed by western blot with rat anti-HA high affinity monoclonal antibodies (Sigma-aldrich, Missouri, United States) at 1:1000 dilution for KPC-2 and its variants. Goat anti-rat immunoglobulin Horseradish Peroxidase (HRP) (Invitrogen), at 1:10000 dilution, were used as secondary antibodies. SuperSignal West Pico PLUS Chemoluminescent substrate (ThermoFisher) was use for detection. Protein band intensities were quantified with ImageJ. The Precision Plus Protein Standards Kaleidoscope (BioRad, California, United States) provided molecular weight standards.

### Proteomic analysis

Protein identification and characterization by Liquid Chromatography with tandem mass spectrometry (LC-MS/MS) were carried out at the CBM-CSIC protein chemistry facility (Madrid, Spain). Briefly, in-gel digestion was performed after SDS-PAGE of the periplasmic extract. Gel bands of the target molecular weight were dried, destained in acetonitrile (1:1), reduced with 10 mM DTT at 56°C for 1 hour, and alkylated with 10 mM iodoacetamide for 30 minutes at room temperature in darkness. The proteins were then digested in situ with sequencing-grade trypsin (Promega, Madison, WI), following the method described by Pérez *et al.*^106^. The gel pieces were dehydrated with acetonitrile, then dried in a speedvac. They were rehydrated in 100 mM Tris-HCl (pH 8), 10 mM CaCl2 with 12.5 ng/μl trypsin for 1 hour on ice. After removing the digestion buffer, the gels were incubated with fresh 100 mM Tris-HCl (pH 8), 10 mM CaCl2 at 37°C for 12 hours. Digestion was stopped by adding 1% TFA. The supernatants were dried and desalted using ZipTip C18 pipette tips (Millipore) before mass spectrometric analysis.

The desalted protein digest was resuspended in 10 µl of 0.1% formic acid and analyzed by Reverse Phase Liquid Chromatography coupled with Tandem Mass Spectrometry (RP-LC-MS/MS) using an Easy-nLC II system and an LTQ-Orbitrap-Velos-Pro mass spectrometer (Thermo Scientific). Peptides were concentrated on-line with a 0.1 mm × 20 mm C18 RP precolumn (Thermo Scientific) and separated using a 0.075 mm × 250 mm bioZen 2.6 µm Peptide XB-C18 RP column (Phenomenex) at 0.25 μl/min. A 60-minute dual gradient was applied: 5−25% solvent B for 45 min, 25−40% solvent B for 45 min, 40−100% solvent B for 2 min, and 100% solvent B for 18 min (Solvent A: 0.1% formic acid in water; Solvent B: 0.1% formic acid, 80% acetonitrile in water). ESI ionization was performed with a Nano-bore Stainless Steel emitter (Proxeon) at 2.1 kV, with S-Lens set at 60%. The Orbitrap resolution was 30,000. Survey scans (400–1600 amu) were followed by twenty data-dependent MS/MS scans (Top 20), with an isolation width of 2 u, normalized collision energy of 35%, and dynamic exclusion for 60 seconds. Charge-state screening rejected unassigned and singly charged ions.

Peptide identification from raw data was carried out using PEAKS Studio vXPro/ v11.5 search engine (Bioinformatics Solutions Inc., Waterloo, Ontario, Canada). Database search was performed against uniport-escherichia-coli.fasta (4403 entries; UniProt release 05/2023). (decoy-fusion database). The sequence of the KPC-2-HA protein has been included in this database. The following constraints were used for the searches: tryptic cleavage after Arg and Lys, up to two missed cleavage sites, and tolerances of 20 ppm for precursor ions and 0.6 Da for MS/MS fragment ions and the searches were performed allowing optional Met oxidation and Cys carbamidomethylation. False discovery rates (FDR) for peptide spectrum matches (PSM) and for protein was limited to 0.01. Only those proteins with at least two unique peptides being discovered from LC/MS/MS analyses were considered reliably identified.

### Statistical analyses and data visualization

All statistical analyses were conducted using the R programming language (version 3.6.3), utilizing built-in functions and available packages. Pearson correlation coefficients, linear regression, Student’s t-Test, Wilcoxon Rank Sum test, and pairwise Wilcoxon signed-rank tests were performed with the functions *cor(), lm(), t.test(), wilcox.test()*, and *pairwise.wilcox.test()*, respectively. Data visualization for Figures 1C and 2C used the “pheatmap” and “rgl” libraries, respectively. Figure 3A was generated using ChimeraX.

## ETHICS

This work did not require ethical approval from a human subject or animal welfare committee.

## DATA ACCESSIBILITY

All data necessary to replicate the research findings is available, either at public data repositories (TBD) or in the electronic supplementary material (TBD).

## AUTHORS’ CONTRIBUTIONS

**LD:** Conceptualization (equal); methodology (lead); investigation (lead); formal analysis (supporting); writing – original draft (supporting); writing – review and editing (equal), funding acquisition (supporting), supervision (supporting). **IN**: investigation (supporting); writing – review and editing (supporting). **AC:** Conceptualization (equal); formal analysis (lead); writing – original draft (lead); writing – review and editing (equal), funding acquisition (lead), supervision (lead).

## ACKNOWLEDGMENTS

We thank J. Barber and L. López-Merino for proofreading the manuscript.

---

**Figure S1.**
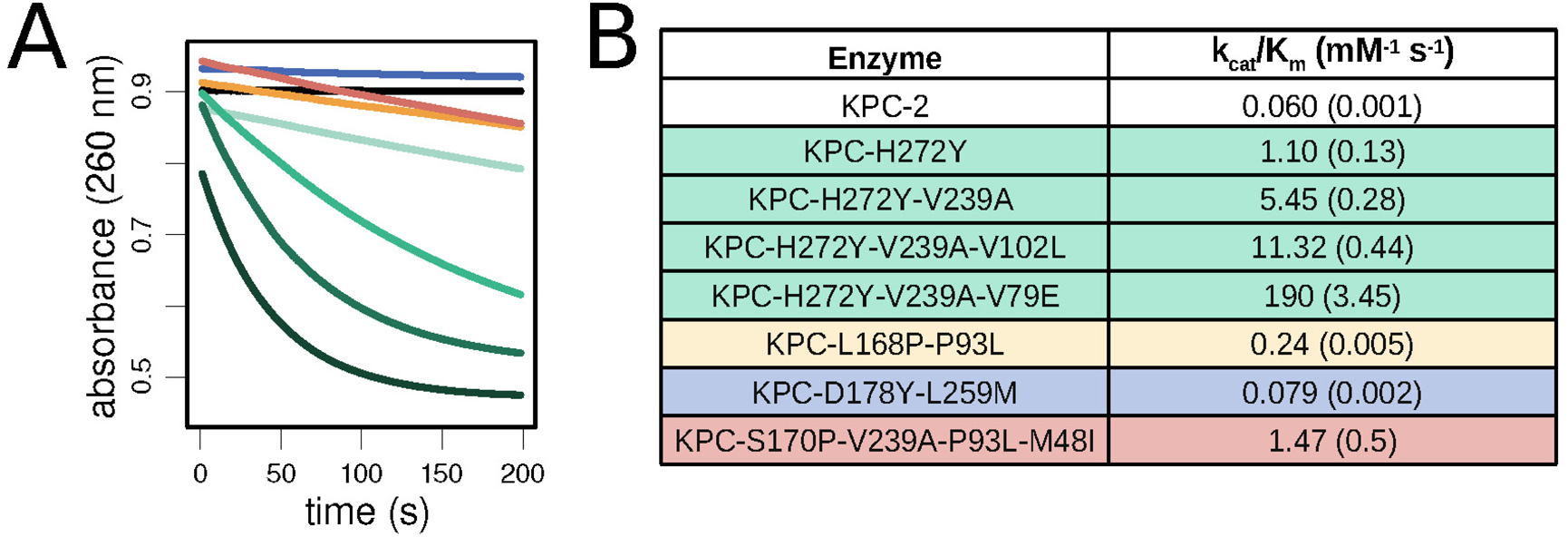
Progress curves of CAZ hydrolysis from a colorimetric assay. All reactions were performed with 500 nM enzyme and 50 μM ceftazidime. Hydrolysis of ceftazidime results in a decrease in absorbance at 260 nm. Curves correspond to: KPC-2 (black), KPC-D178P-L260M (blue), KPC-L168P-P94L (yellow), KPC-S170P-V240A-P94L-M49I (orange), KPC-H272Y (very light green), KPC-H272Y-V240A (light green), KPC-H272Y-V240A-V103L (green), and KPC-H272Y-V240A-V80E (dark green). **(B)** Steady-state kinetic parameters for CAZ hydrolysis for ancestral KPC-2 and its variants.

**Figure S2.**
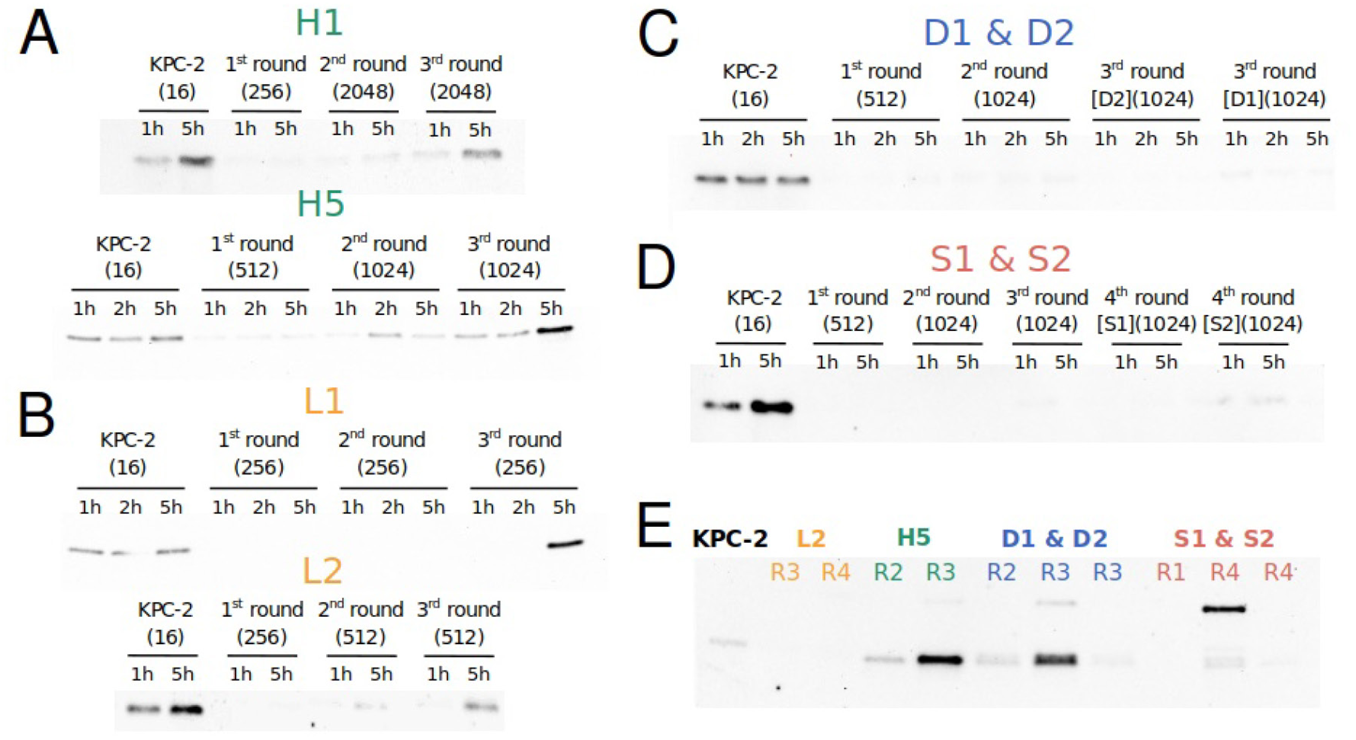
Western blot of KPC-2 and its variants in periplasmic extracts of *E. coli* TOP10. (A-D) Periplasmic abundance of representative lineages from groups H272 (green), L168 (yellow), D178 (blue), and S170 (red) after 1 and 5 hours of growth. **(E)** Immunoblot of selected KPC variants in the periplasm of *E. coli* TOP10 after 10 hours of incubation. Colors as above.

**Figure S3.**
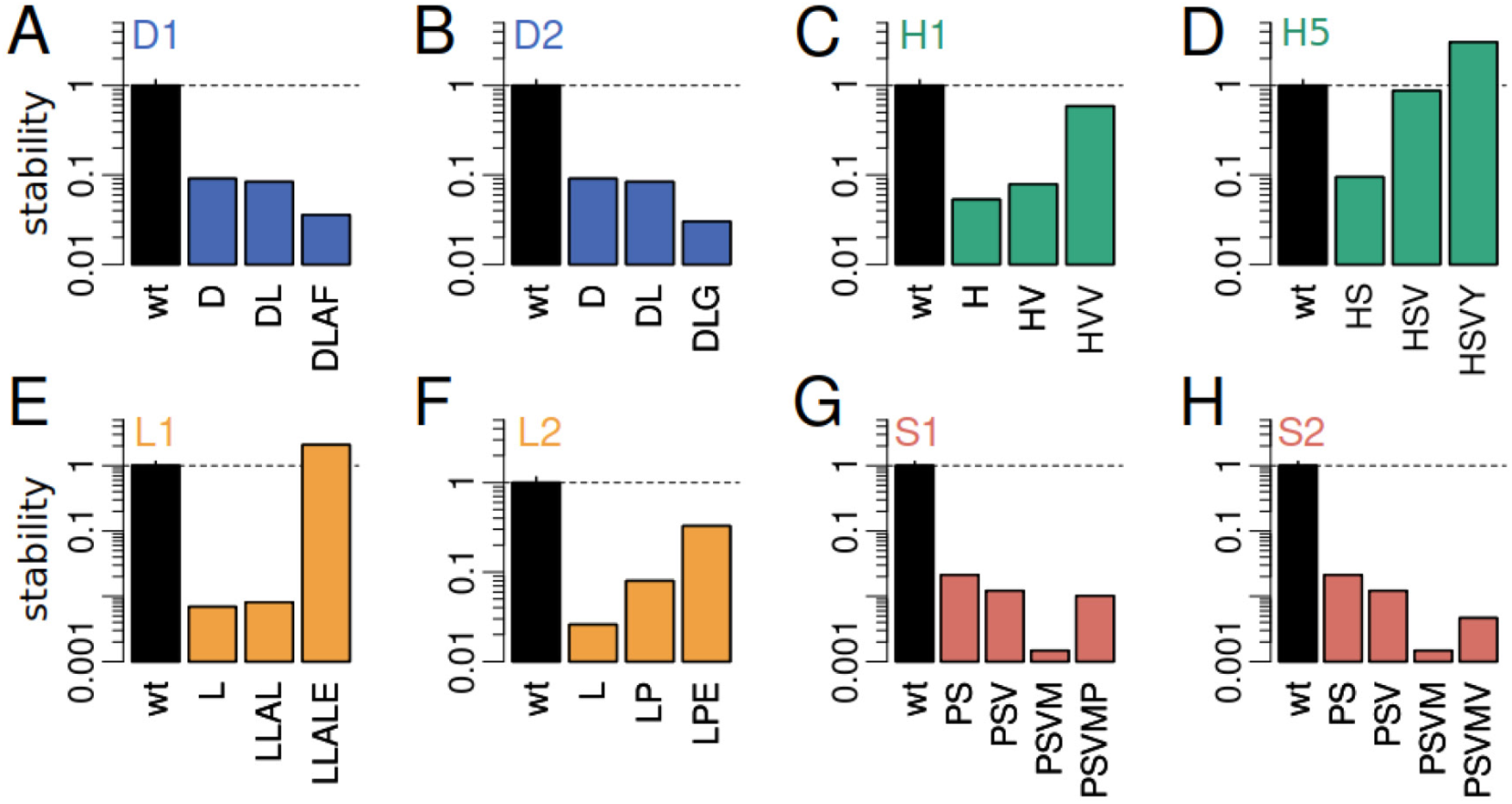
Periplasmic stability of selected variants. Bar plots show the log intensity ratio of initial (1 h) and final (5 h) bands for each variant, quantified from the Western blots in Figure S2. Lineage labels are in the upper left. Colors as in Figure S2.

**Figure S4.**
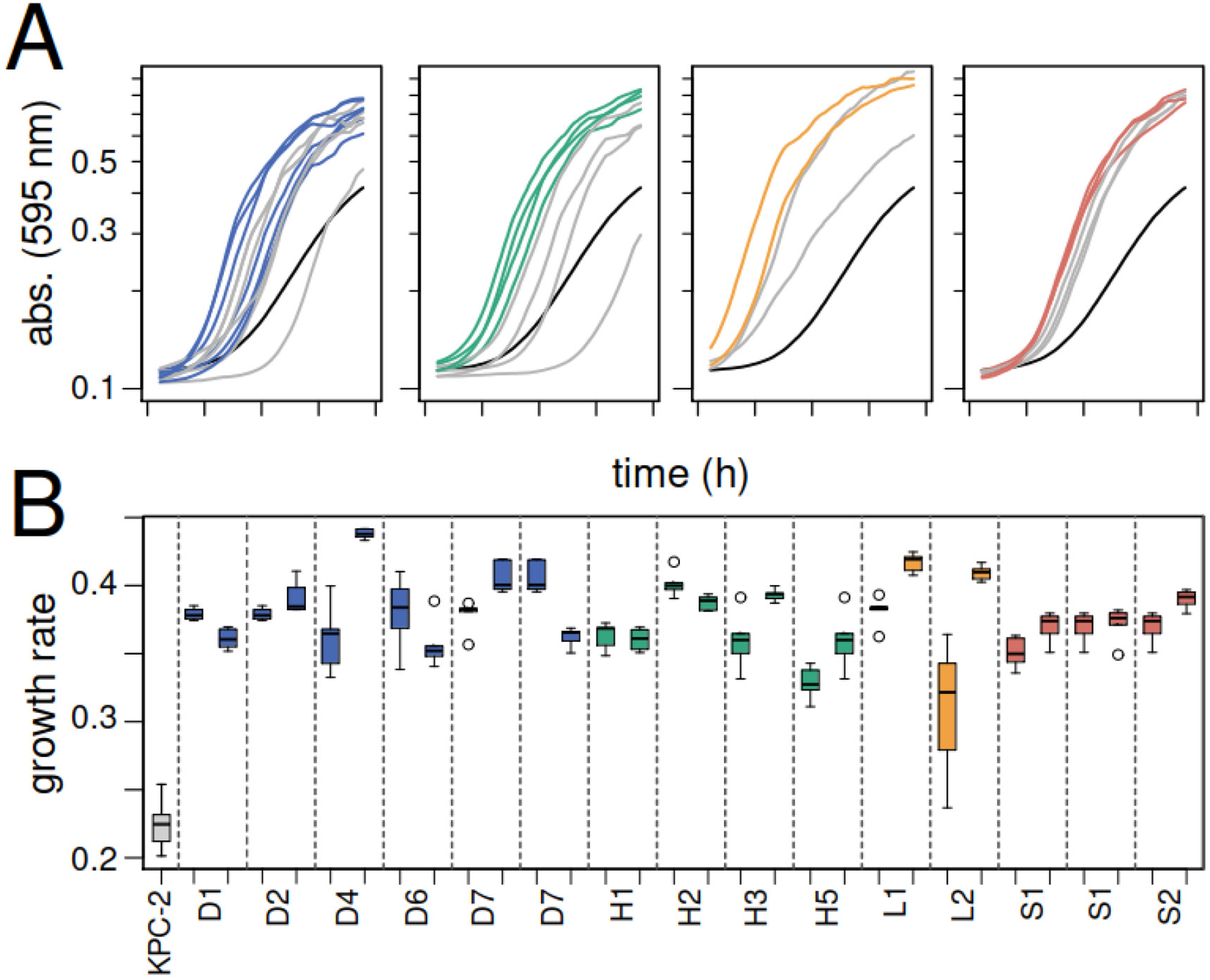
Growth rates of consecutive mutants within plateaus. **(A)** Optical density changes over 5 hours of growth. The black line represents the strain with wild-type KPC-2, gray lines represent earlier plateau strains, and colored lines represent subsequent strains. **(B)** Boxplot summarizing maximum growth rates from the growth curves above.

**Table S1.**
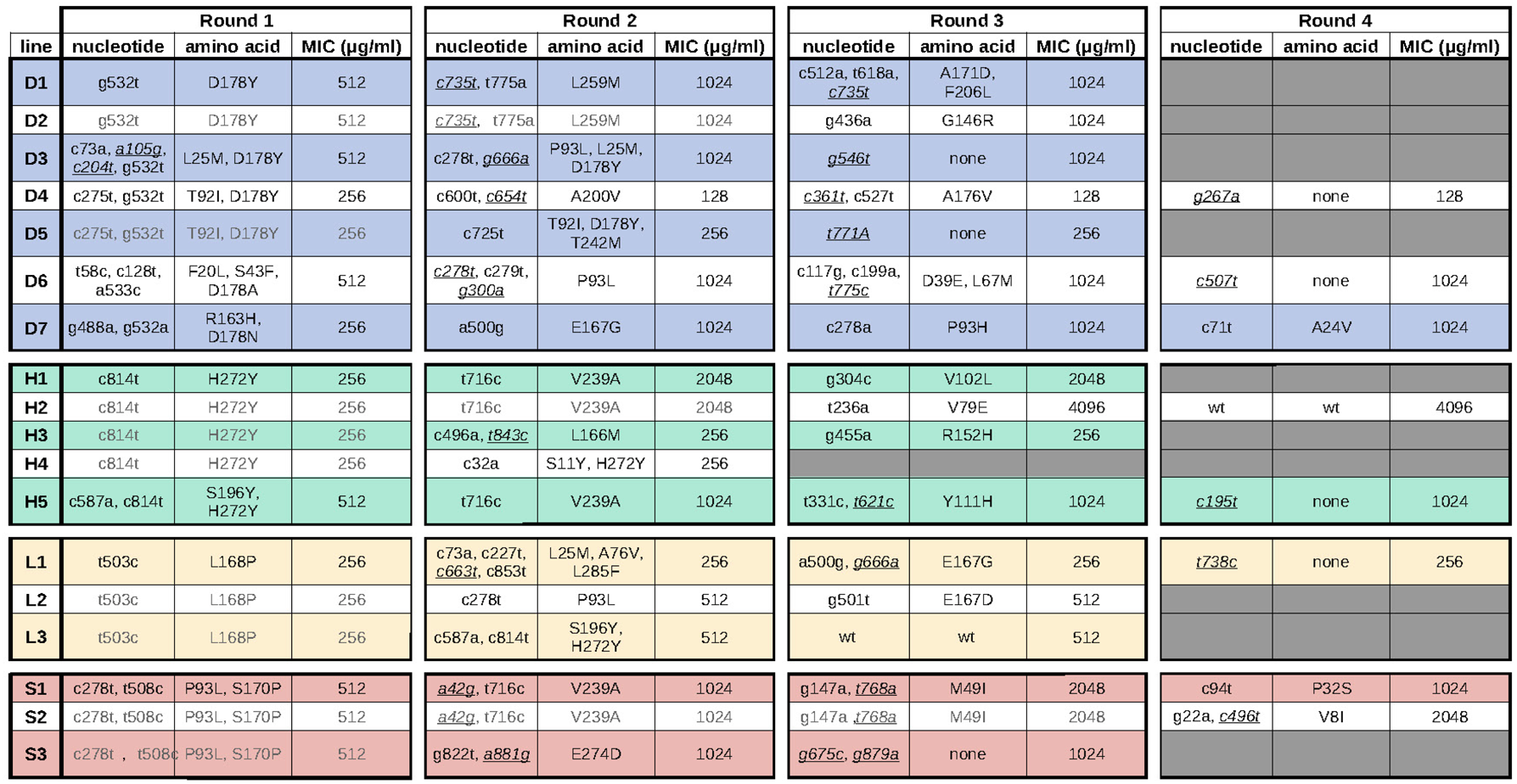
Mutations recovered across different rounds of directed evolution. Lineages are grouped by the presumed first beneficial mutation, color-coded as blue (D178Y), green (H272Y), yellow (L168P), and red (S170P). Each round includes three columns showing changes at the nucleotide level in the *blaKPC* gene, at the amino acid level in the KPC protein, and MIC values for CAZ. Synonymous mutations are indicated by underlined, italicized nucleotide changes.

